# A size-invariant bud-duration timer enables robustness in yeast cell size control

**DOI:** 10.1101/211714

**Authors:** Corey A.H. Allard, Franziska Decker, Orion D. Weiner, Jared E. Toettcher, Brian R. Graziano

## Abstract

Cell size drives key aspects of cell physiology, including organelle abundance [1, 2] and DNA ploidy [3]. While cells employ diverse strategies to regulate size [4–11], it is unclear how they are integrated to provide robust, systems-level control. In budding yeast, a molecular size sensor restricts passage of small cells through G1, enabling them to gain proportionally more volume than larger cells before progressing to Start [7, 12, 13]. Size control post-Start is less clear. S/G2/M duration in wildtype cells shows only a weak dependence on cell size; and since yeast exhibit exponential growth, larger cells would be expected to add more volume than smaller ones [7, 14–17]. However, even large mother cells produce smaller daughters, suggesting that additional regulation may occur during S/G2/M [7]. To gain further insight into post-Start size control, we prepared ‘giant’ yeast (>10-fold larger than typical volume) using two approaches to reversibly block cell cycle progression but not growth: optogenetic disruption of the cell polarity factor Bem1 [18, 19] and a temperature-sensitive *cdk1* allele [20]. We reasoned that giant yeast would satisfy pre-Start size control while enabling us to uncover post-Start size-limiting mechanisms though the identification of invariant growth parameters. Upon release from their block, giant mothers reenter the cell cycle and their progeny rapidly return to the original unperturbed size. This behavior is consistent with a size-invariant ‘timer’ specifying the duration of S/G2/M and indicates that yeast use at least two distinct mechanisms at different cell cycle phases to ensure size homeostasis.

## RESULTS AND DISCUSSION

### Preparing ‘giant yeast’ *via* isotropic cell growth

To achieve reversible control over cell size in the budding yeast *S. cerevisiae*, we first took advantage of the light-responsive PhyB/PIF optogenetic system [21] to control the localization of Bem1, a cell polarity factor [18, 19]. In this “optoBem1” system, red light illumination re-localizes the PIF-Bem1 fusion protein to mitochondria-anchored PhyB (**Fig. 1A**). Light-induced Bem1 re-localization produces an acute loss-of-function phenotype where cells fail to form a site of polarized Cdc42 activity, fail to initiate budding, and instead undergo continuous isotropic growth [18] (**Fig. 1B-C**). Strikingly, this effect is quickly reversed upon illumination with infrared (IR) light, which releases PIF-Bem1 from the mitochondria within seconds. Upon release, cells form a bud within minutes and proceed to cytokinesis (**Fig. 1, B and D**). The PIF-Bem1 fusion protein appears to fully recapitulate normal Bem1 function: when it is not sequestered to the mitochondria, overall cell sizes and cell growth rates are similar to an isogenic wildtype strain [18].

**Figure 1.**
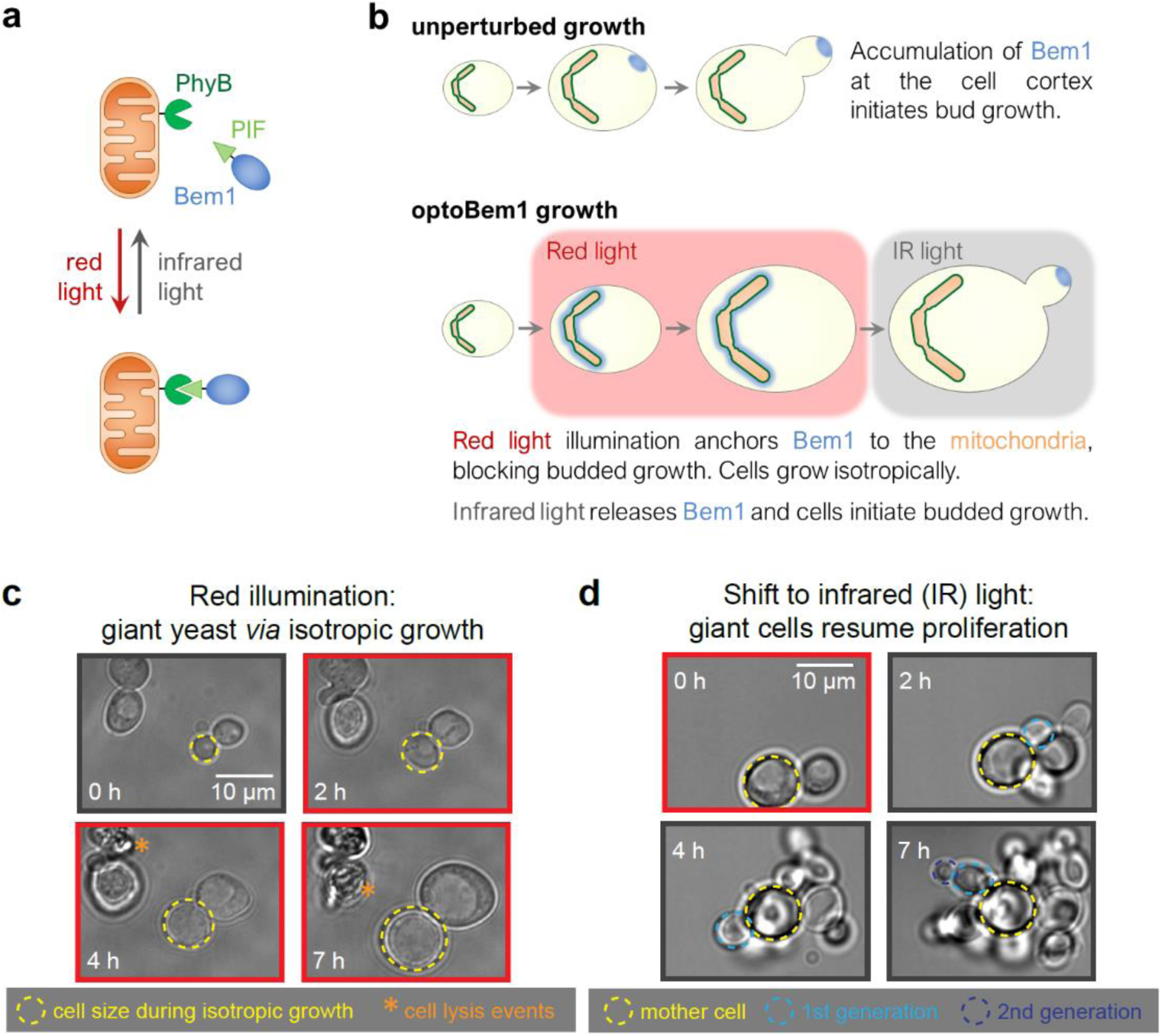
Control of yeast cell size using optogenetics. **(a)** Exogenous PhyB (phytochrome B; dark green) is fused to the C-terminus of S. cerevisiae Tom20, anchoring it to the mitochondria outer membrane (orange). PIF (phytochrome-interacting factor; light green) is fused to endogenous Bem1 (blue). Illumination with red light drives a conformational change in PhyB allowing it to bind PIF-Bem1. Conversely, illumination with infrared (IR) light drives the reverse reaction, releasing PIF-Bem1. **(b)** Production of ‘giant’ yeast using reversible optogenetic-based Bem1 disruption. **(c)** Cells were illuminated with red light for 8 h (indicated by red borders) and imaged every 10 min using bright-field microscopy. **(d)** Following red light illumination for 6-10 hours as in **1C**, large cells were illuminated with IR light (indicated by grey borders) and imaged every 5-10 min using bright-field microscopy.

We performed additional experiments to more completely characterize optoBem1 giant cells. Our initial experiments quantifying the growth of red light-illuminated optoBem1 cells revealed two subpopulations of cells that grew at different rates (**Fig. S1A**). We hypothesized that cell growth rates differed depending on the cell cycle phase at the time of Bem1 disruption. Indeed, we found that synchronizing optoBem1 cells before red light stimulation led to unimodally-distributed growth (**Fig. S1B-C**). Furthermore, restricting our analysis to measure growth only following entry into G1 yielded a unimodal distribution (**Fig. S1D-H**). We also observed that a substantial fraction of optoBem1 yeast burst as they become increasingly large (**Fig. 1C**, asterisks; [18]), and hypothesized that cell lysis may be a result of large cells’ increased susceptibility to osmotic pressure. Supporting this hypothesis, growing cells in high-osmolarity media containing 1 M sorbitol decreased the frequency of cell lysis (**Fig. 2A**) without affecting the rate of isotropic growth (**Figs. 2B and S1A**). We therefore supplemented our media with sorbitol for all subsequent experiments involving optoBem1-arrested cells. Finally, to test whether growth was isotropic during the entire time period, we pulsed cells with fluorescent Concanavalin A (FITC-ConA) to mark the existing cell wall, followed by a washout of free FITC-ConA. We found that cells exhibited uniform dilution of FITC-ConA around their surface, consistent with isotropic growth **(Fig. 2C)**.

**Figure 2.**
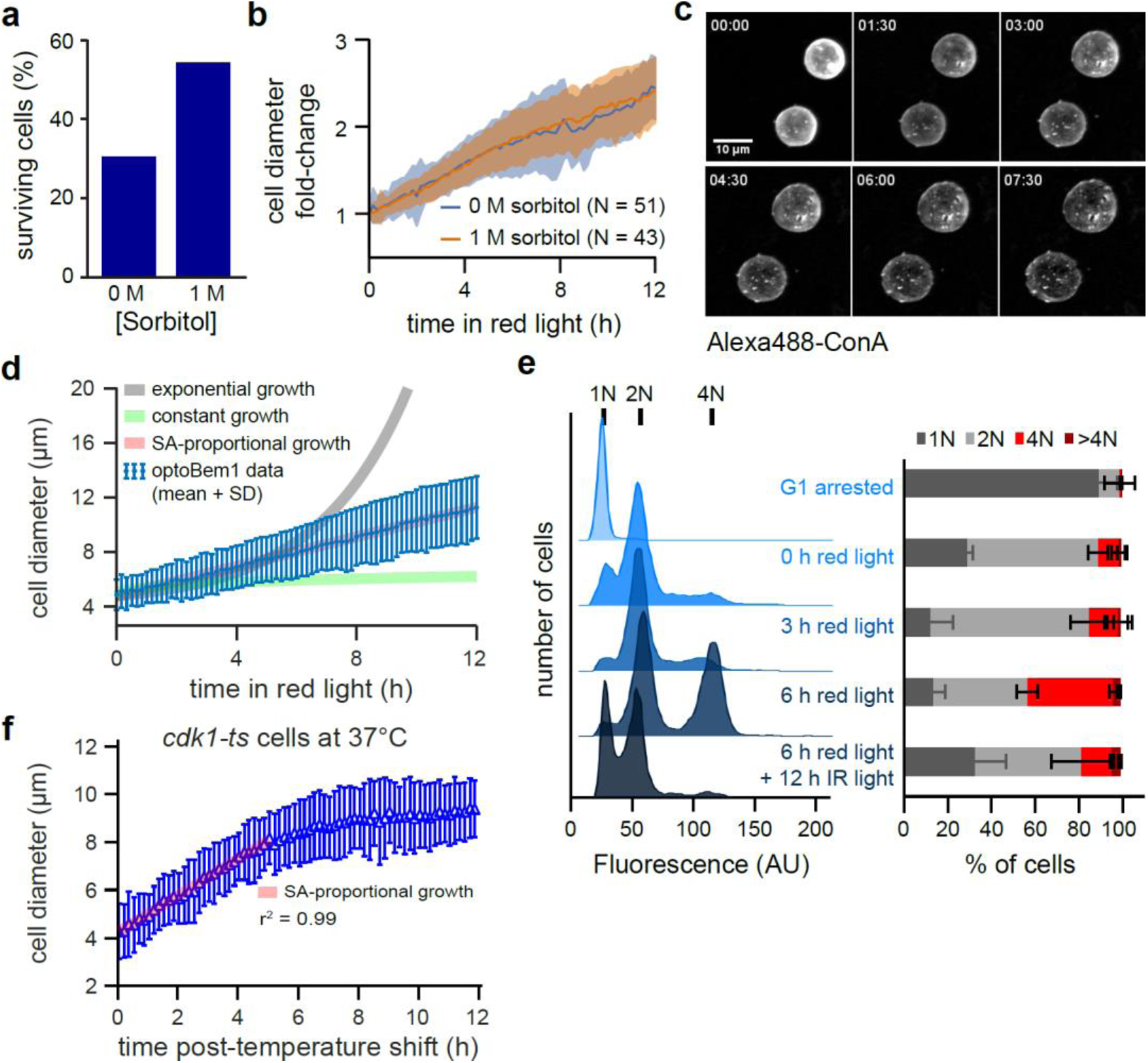
Effects of continuous isotropic growth on yeast physiology. **(a)** Yeast were prepared and imaged as in **Fig. 1C-D** with media containing indicated concentrations of sorbitol. Each bar indicates the percentage of cells surviving the entire 12-h time-course. **(b)** Average normalized diameter of yeast grown in media with or without 1 M sorbitol. N > 40 cells for each condition. Error bars, SD. **(c)** Cell wall staining. Cells were prepared as in **2A**, treated with 100 µg/mL Alexa488-labeled concanavalin-A, and imaged every 15 min for 8 h. Each panel is a maximum intensity projection of multiple focal planes acquired via confocal microscopy. Yeast grow isotropically following Bem1 deactivation. **(d)** Average growth trajectory for Bem1-arrested cells calculated as in **2B** for yeast in 1 M sorbitol from experiments described in **2A**. Each blue dot represents an average of 48 cells. Expected trajectories for exponential, constant, and surface-area-proportional growth are indicated by grey, green, and red lines, respectively. **(e)** Analysis of Bem1-disrupted cells by flow cytometry. Representative plots, left; averaged data from 2 independent experiments, right. Error bars, SD. **(f)** Average growth trajectory for *cdk1-ts* cells at 37 °C. The relationship between diameter and time remained linear for the first 5 h of growth (r^2^ = 0.998), but growth rapidly stalled at later timepoints. N = 126 cells.

### In budding yeast, the rate of isotropic growth during G1 is proportional to cell surface area

Prior studies have established that unperturbed, freely-cycling budding yeast cells appear to exhibit an exponential growth in volume over time [14, 16, 17, 22]. However, most of this growth is localized to the bud, with only a minor contribution from the mother cell’s isotropic growth during G1. The mode of growth may also change depending on cell cycle phase [23]. Since distinguishing between growth patterns is difficult to achieve during the growth interval of normal sized yeast [24], we reasoned that the ability to prepare isotropically-growing yeast with volumes spanning an order of magnitude would permit high-quality measurements of this growth law, and potentially reveal processes that limit cell growth as size increases.

We imaged optoBem1 cells during red light illumination at multiple z-planes and used a custom code to automatically measure cell diameter every 10 min over a 12 h period. Following entry into G1 after Bem1 arrest, we found that isotropically-growing optoBem1 cells exhibited a linear increase in cell diameter over time, corresponding to a rate of volume growth proportional to *(time)*^*3*^ (**Fig. 2D**). Since these volume increases also show a strong correlation with protein content, as assessed by fluorescence [14] (**Fig. S1I**), our data suggest that the growth we observed primarily arises from increases in cell mass rather than cell swelling (e.g., water influx). This result is inconsistent with two classic models of cell growth: a constant growth law, where volume increases linearly over time; and exponential growth, where the rate of growth is proportional to the cell’s current volume. In contrast, a linear increase in cell diameter is the expected result for volume increasing in proportion to cell surface area (Supplemental Experimental Procedures). Surface area-proportional growth could arise if nutrient/waste exchange across the plasma membrane is a limiting factor for growth [25].

We observed that red light-illuminated optoBem1 cells also exhibited a change in DNA content over time. While most cells maintained a ploidy of 2N or less during the first 3 h of Bem1 disruption, a population of 4N cells appeared following 6 h of arrest (**Fig. 2E**), consistent with prior reports suggesting that after Bem1 disruption, some arrested cells eventually leak through the cell cycle block and undergo DNA endoreduplication [26]. To ensure that the surface area-proportional growth was not an artifact of increased ploidy, we set out to generate ‘giant yeast’ *via* a second, non-optogenetic method: disruption of Cdk1/Cdc28 using the temperature-sensitive allele *cdc28-13* (hereafter referred to as *cdk1-ts*). Unlike optoBem1 cells, nearly all *cdk1-ts* cells at the restrictive temperature arrest in G1 without undergoing further DNA replication [20]. We found that *cdk1-ts* cells grown at the restrictive temperature to induce arrest in G1 also exhibited a linear increase in cell diameter, consistent with growth proportional to surface area (**Fig. 2F**). However, *cdk1-ts* were unable to maintain this rate of growth over the entire 12-h time-course: After reaching a volume of 500-700 μm^3^ (6-7 h following the temperature shift), cell growth stalled (**Fig. S1J**).

Taken together, our results from both optoBem1 and *cdk1-ts* cells indicate that the isotropic growth rate during G1 is proportional to surface area over a wide range of cell sizes. DNA endoreduplication does not appear to affect this overall growth rate but may be required to sustain it beyond a critical cell size, giving rise to the robust continued growth of optoBem1 cells. It has been shown in other organisms, for example, that DNA endoreduplication enables large increases in cell size [27]. Our findings can be reconciled with prior observations of exponential growth in wildtype budding yeast in at least two ways. First, exponential growth might only govern bud growth, masking a distinct growth law that operates in isotropically-growing cells. Second, it is possible that cells become surface area-limited at sizes just above that of wildtype cells, thereby inducing a shift from volume-proportional growth to surface area-proportional growth.

### Giant yeast retain size homeostasis

Cell size control pathways exist to correct for deviations from a set-point size, yet most previously-identified size control pathways specifically operate on cells that are born too small, delaying cell cycle progression to enable further growth to occur [6, 28]. Because the light and temperature-shift stimuli with which we prepared ‘giant’ yeast are fully reversible, we reasoned that we could monitor the return to a steady-state size distribution after releasing giant cells from their block.

We prepared giant optoBem1 cells by incubating them in red light for 8 h and monitored them by live-cell microscopy after releasing them into infrared light. Strikingly, we found that cell populations rapidly returned to their unperturbed state (**Fig. 3A**), with individual daughter cells reaching the set-point volume in as few as three rounds of division (**Fig. S2A**). Return to the set-point size is not driven by cell shrinking, as giant mothers maintained their maximum volume over multiple rounds of budding (e.g., **Fig. 1D**). Instead, the giant mothers are eventually diluted out as successive generations are born, an effect that is especially prominent in cell populations at least 10 h post-Bem1 release (**Fig. 3A**). In these populations, size distributions have a single mode near the set-point volume but exhibit long tails towards larger volumes (**Fig. S2B**). Our observation that cell size recovers after only a few generations strongly supports the existence of size control acting on large cells and demonstrates that size homeostasis across a cell population is robust even to extreme increases in cell volume.

**Figure 3.**
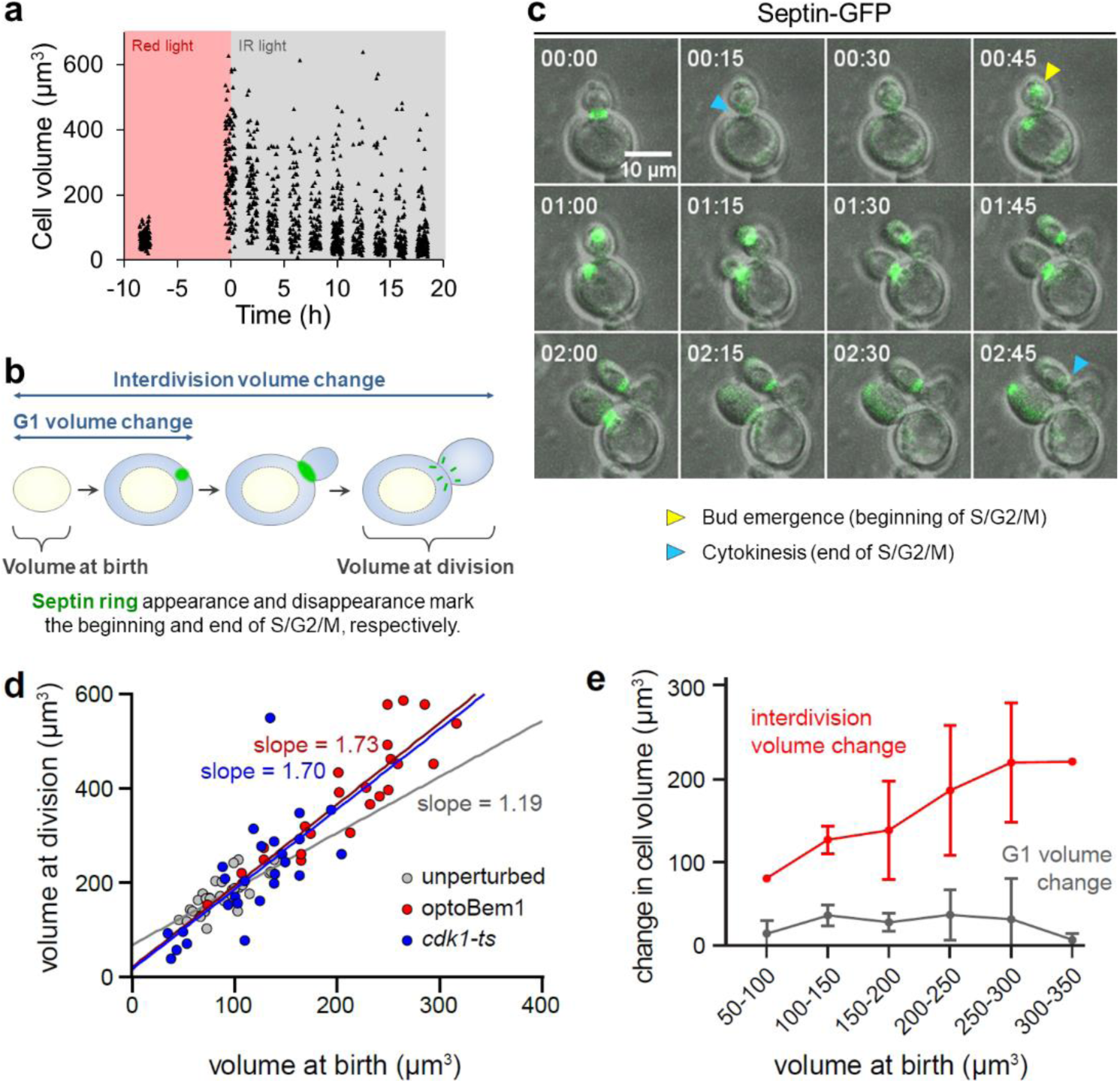
Convergence of yeast to set-point volume is inconsistent with an ‘adder’. **(a)** Cells were incubated under red light illumination for 8 h followed by IR light illumination for 18 h. At indicated timepoints (every 2 h during IR light illumination), cell volumes were measured by microscopy. Each point represents a single cell. **(b)** Budding yeast cell cycle with labels depicting volume and growth intervals measured in **3C-E**. Blue-shaded areas, volume added as a newly-born cell grows. **(c)** OptoBem1 cells were illuminated for 8-10 h with red light (to generate giant yeast), then switched to IR light (allowing giant yeast to bud and divide) and imaged every 5 min for ∼8 h. *cdk1-ts* cells were incubated at 37 °C for 8 h, then switched to 25 °C prior to imaging. Exogenously-expressed Cdc10-GFP was used to mark septin rings (green). Panels depict representative optoBem1 cells. Time, HH:MM. **(d)** Each point represents a single cell. Colored lines, best-fit line by linear regression analysis. **(e)** Cells were binned by volume at birth, as indicated. N = 28 optoBem1 cells.

### Convergence of yeast to a set-point size does not rely on adder-based mechanisms

Quantitatively monitoring cell growth in yeast—as well bacterial, archaeal, and mammalian cells— has shown that the behavior of many organisms is consistent with an adder that monitors size across an entire cell cycle to correct for deviations in cell size and maintain size homeostasis in the population [10, 17, 29, 30]. However, a recent study argued that in budding yeast, the adder behavior could arise from independent regulation of pre- and post-Start events, without a cell needing to keep track of its added volume across all cell cycle phases, and may fail under various perturbations [31]. To test whether adder-based mechanisms could account for size control in giant yeast, we quantified interdivision volume change in successive cell division cycles after releasing optoBem1 cells into infrared light. For this experiment we prepared optoBem1 cells that also expressed fluorescently-labeled septin rings [32], which enabled us to time both bud emergence and cytokinesis and thus separate pre-Start and post-Start size regulation (**Fig. 3B-C**; see Methods).

The ‘adder’ model predicts that the cell volume at division (*v*_*d*_, which includes both the mother and bud compartments) should be proportional to cell volume at birth (*vb*) with a slope of 1 (i.e. *vd= v*_*b*_ *+* δ, where δ represents the constant volume increment ‘added’ through one cell cycle (**Fig. 3B**, rightmost blue-shaded area)) [17]. Indeed, for unperturbed cells, we found that cell volume at division was linearly related to volume at birth with a slope of 1.19 (95% CI, 0.82-1.56) (**Fig. 3D**). However, we found that the adder model poorly explained the cell size relationships in our giant cells, where the volume at division was related to volume at birth with a slope of 1.73 (95% CI, 1.32-2.13) (**Fig. 3D**). This relationship was also evident when individual cells were tracked over time: the interdivision volume change, δ, was positively correlated with the volume at birth (**Fig. 3E**). This size-dependent volume change occurred entirely during S/G2/M phase, as cells added a minimal volume during G1 that did not vary with cell size (**Fig. 3E**, grey curve). We also performed analogous experiments in *cdk1-ts* giant cells that were shifted back to the permissive temperature. These experiments revealed a similar relationship: large cells grew more than small cells, exhibiting a linear relationship between volume at division and volume at birth with a slope of 1.70 (95% CI, 0.96-2.44) (**Fig. 3D**). These results are broadly consistent with recent work showing that although size control in unperturbed cells resembles an adder-based mechanism, no mechanistic adder regulates volume addition across the entire cell cycle [31]. Our data also suggest that any size regulation limiting the growth of large cells is likely a consequence of regulation in S/G2/M, as growth during G1 is negligible.

### A ‘timer’ specifying budding duration operates across a broad range of cell sizes

If an adder is unable to explain size homeostasis in giant yeast, what regulatory mechanisms or growth laws might operate on the daughters of giant cells during S/G2/M? Two possibilities include a bud ‘sizer’, where bud growth would be restricted after reaching a critical size; and a bud ‘timer’ in which cytokinesis would occur at a fixed duration following the beginning of S/G2/M (i.e. bud emergence) (**Fig. 4A**). Such ‘sizers’ and ‘timers’ have been proposed to operate in a variety of biological systems [24, 33–35]. To distinguish between these possibilities, we tracked the timing of bud emergence and cytokinesis by septin ring appearance and disappearance, respectively, following reactivation of Bem1 in giant optoBem1 cells (**Fig. S3A**). Daughter volume strongly correlated with mother volume (**Fig. 4B**), inconsistent with a bud sizer mechanism. Our prior observation that the interdivision volume change scales positively with cell birth size (**Fig. 3E**) further argues against a bud sizer for cell volume control. In contrast, our data were consistent with a timer specifying the duration of S/G2/M: the time from bud emergence to cytokinesis did not vary as a function of mother cell volume and took average 95 min across cells of all volumes (**Fig. 4C**).

**Figure 4.**
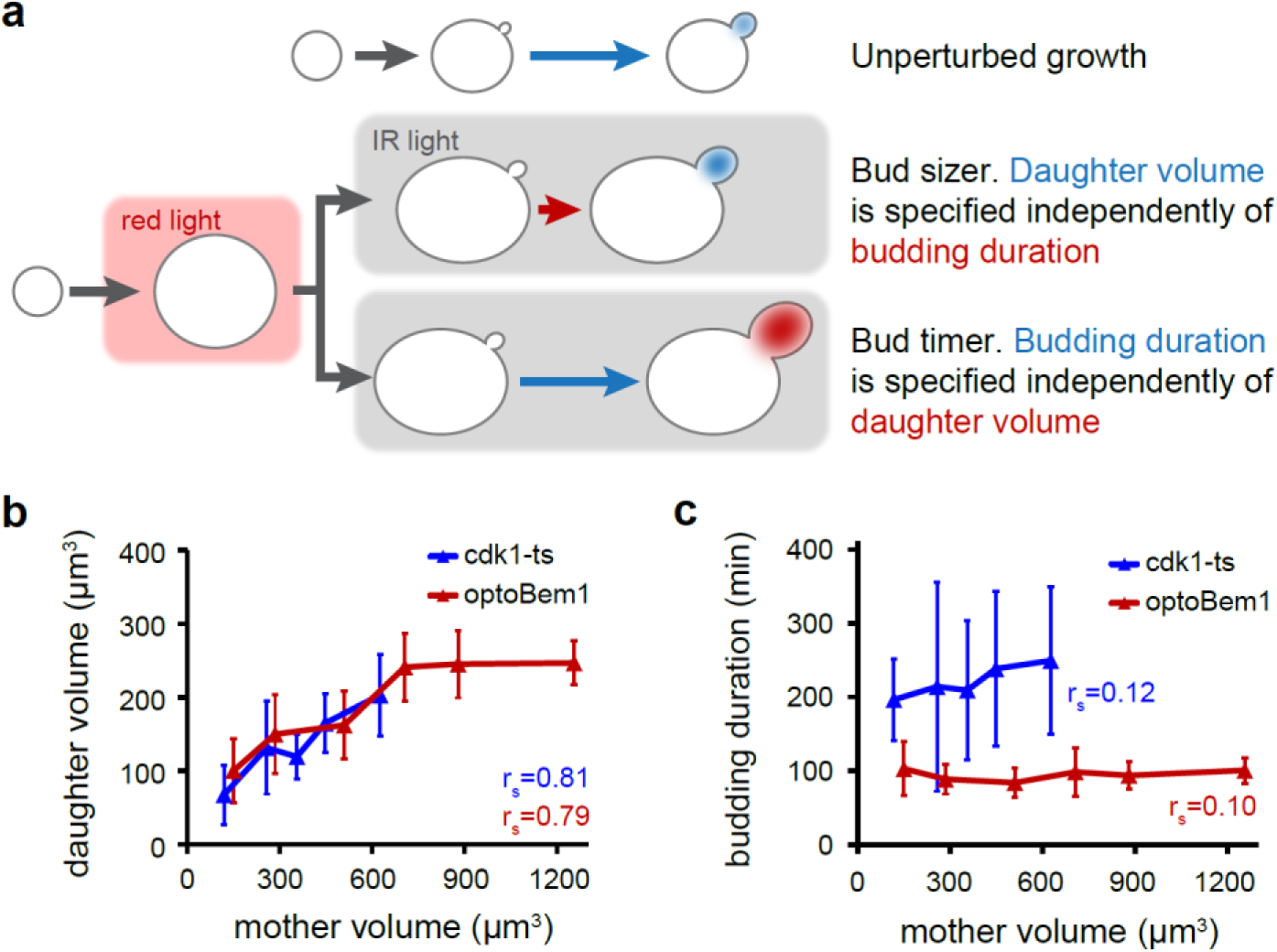
Comparison of sizer and timer mechanisms for regulation of bud growth. **(a)** Schematic depicting the use of our optogenetics system for discriminating between bud ‘sizer’ and bud ‘timer’ mechanisms for specifying daughter cell volume. **(b and c)** Cells were binned by mother volume in 200-µm^3^ increments. The average volume within each bin is plotted. Measurements of ‘budding duration’, ‘mother volume’, and daughter volume’ were obtained as described in **Fig. S3A**. N = 73 optoBem1 cells and 80 *cdk1-ts* cells, with each bin containing at least 5 cells. Error bars, SD. rs, Spearman’s rho.

Similar experiments performed using *cdk1-ts* cells (**Fig. 4B-C**) were consistent with our observations in optoBem1 cells, revealing a size-independent duration of budding. However, we observed one notable difference: the duration of the size-invariant bud timer in giant *cdk1-ts* cells was substantially longer (215 min) than that of giant optoBem1 cells (95 min) (**Fig. 4C**). As Cdk1 is a key driver of mitosis in eukaryotes [20], the increased duration of the bud timer in *cdk1-ts* cells may arise from the need to refold or synthesize new Cdk1 molecules to complete S/G2/M following a shift from the restrictive to permissive temperature. Furthermore, even when grown at the permissive temperature, the doubling time of *cdk1-ts* cells is longer than an isogenic wildtype strain (**Fig. S3B**), suggesting that *cdk1-ts* may not be able to fully complement *CDK1*.

In summary, we find that a timer specifying a constant budding duration describes how a cell population founded by ‘giant’ cells returns to their set-point volume. Although mother and daughter sizes are correlated across a broad size range, daughters are always born smaller than mother cells. After cytokinesis, daughter cells remaining larger than the set-point volume exhibit a G1 phase with virtually no growth (**Fig. 3E**) and bud rapidly, leading to a geometric shrinking in successive generations (**Fig. 3A**). Indeed, a back-of-the-envelope calculation demonstrates that if newly-budded daughters are each 50% smaller than their mothers, a 32-fold decrease in cell

volume can be achieved in 5 generations (2^5^ = 32). Assuming a 100 min doubling time (**Fig. 4C**), a return to the set-point size would take ∼8 h. A fixed budding time, even in the absence of active molecular size sensors in S/G2/M, is sufficient to buffer against persistence of abnormally large cell sizes in the population. We also note that the bud duration timer we describe is quite complementary to G1-phase size sensors such as Whi5 [13], which compensate for a small size at birth by elongating G1 phase.

Our conclusions are derived from cells prepared using two independent perturbations: optogenetic inactivation of the Bem1 polarity factor and a temperature-sensitive *cdk1* allele. Importantly, each of these perturbations targets distinct cellular processes and thus produces distinct physiological defects. Cells lacking Bem1 activity exhibit weakened cells walls (**Fig. 1C**) and undergo successive rounds of DNA endoreduplication following their initial arrest in G1 (**Fig. 2E**). In contrast, loss of Cdk1 does not produce such defects but its disruption requires incubating cells at 37 °C, which may broadly activate environmental stress response pathways. Furthermore, *cdk1-ts* may not fully complement *CDK1*, even at the permissive temperature (**Fig. S3B**). That each of these perturbations reveals similar mother-daughter size correlations as well as a size-invariant bud timer strongly supports the generality of our conclusions.

The bud timer we describe here need not be a dedicated biochemical circuit to sense budding duration, compare it to a set-point, and dictate the transition to cytokinesis. Its existence could simply arise due to the time required by independent cellular processes that coincide with bud growth, such as the combined duration of S-phase or mitosis. Nevertheless, one observation suggests more complex regulation: the duration of the size-invariant bud timer is markedly longer in enlarged *cdk1-ts* vs. optoBem1 cells (**Fig. 4C**), yet mother-daughter sizes are nearly identical in these two backgrounds (**Fig. 4B**). These data suggest that the duration of the bud timer may be inter-related to Cdk1 activity and cells’ growth rate during S/G2/M. Recent work has found that mitosis and bud growth rate are closely coordinated and that cells may extend the duration of mitosis to compensate for slow growth that occurs under poor nutrient conditions [36]. Dissecting the dependencies between growth rate, Cdk1 activity and the duration of post-Start events presents a promising direction for future study.

## MATERIALS AND METHODS

### Strains, plasmids, and growth conditions

All yeast strains used are isogenic to an ‘optoBem1’ strain which was created in the w303 genetic background and contained exogenous PhyB-mCherry-Tom7 with endogenous Bem1 C-terminally tagged with mCherry-PIF, as previously described [18]. The *cdc28-13* (*cdk1-*ts) strain was a kind gift from David Morgan. A pACT1-CDC10-eGFP expression vector was created by Gibson assembly [37], with the CDC10 expression cassette inserted between the NotI and XmaI sites of the pRS316 vector [38]. For the experiments described in **Figs. 3-4**, **S1D-G, and S2**, the indicated vector was transformed into our optoBem1 or *cdk1-*ts strain and selection was maintained by growing yeast in synthetic complete media lacking uracil (Sunrise). For all other experiments, yeast were cultured in synthetic complete media (Sunrise).

### Preparation of giant yeast

Preparation of yeast prior to optogenetic experiments was performed, in general, as previously described [18]. Yeast undergoing exponential growth in synthetic media (with or without 1 M sorbitol) were treated with 31.25 µM phycocyanobilin (PCB; Santa Cruz Biotechnology, Inc.) and incubated in foil-wrapped tubes (to block light) at 30 °C for a minimum of 2 h. For all microscopy experiments, yeast were spun onto glass-bottom 96-well plates (Brooks) coated with Concanavalin A and washed once with fresh PCB-containing media (with or without sorbitol) to remove floating cells. Cells remained approximately spherical following this procedure, as assessed by Concanavalin A staining (e.g., **Fig. 2C**). Mineral oil (Sigma-Aldrich) was then carefully layered on top of each sample to prevent evaporation. Imaging was performed at room temperature. For experiments where isotropic growth was measured (e.g., **Fig. 2A**), yeast were plated and imaged immediately following PCB treatment. For experiments where growth following Bem1 reactivation was examined (e.g., **Fig. 3**), PCB-treated yeast were first placed in clear culture tubes and incubated at room temperature for >6 h while undergoing constant illumination with a red LED panel (225 Red LED Indoor Garden Hydroponic Plant Grow Light Panel 14W, HQRP). Cells were then plated and imaged.

For experiments involving the *cdk1-ts* strain, cells were maintained in liquid cultures of synthetic complete media at 25 °C for at least 24 h and plated as described for the optoBem1 strain. Imaging was performed at 37 °C for experiments where isotropic growth during G1 was measured (e.g., **Fig. 2F**). For experiments where size control was assessed (e.g., **Fig. 4B-C**), cells were incubated at 37 °C for 8 hr, then shifted to 25 °C 30 min prior to imaging.

### Microscopy

For isotropic growth experiments, samples were imaged on a Nikon Eclipse Ti inverted microscope equipped with a motorized stage (ASI), a Lamba XL Broad Spectrum Light Source (Sutter), a 60x 1.4 NA Plan Apo objective (Nikon), and a Clara interline CCD camera (Andor). Samples were imaged by bright-field microscopy every 10 min for 12 h. Throughout experiments involving optoBem1 cells, a red LED panel (HQRP) was carefully balanced against the motorized stage and microscope body to provide oblique illumination to the cells and ensure that Bem1 remained deactivated. Generous amounts of lab tape (Fisher) were applied to the LED panel and scope to prevent slippage during image acquisition and stage movement.

For the remaining experiments, samples were imaged on one of two spinning disk confocal microscopes, both of which were Nikon Eclipse Ti inverted microscopes with motorized stages (ASI). The first microscope was equipped with a Diskovery 50-µm pinhole spinning disk (Spectral Applied Research), a laser merge module (LMM5, Spectral Applied Research) with 405, 440, 488, 514, and 561-nm laser lines, a 60x 1.49 NA TIRF Apo objective (Nikon), and a Zyla sCMOS camera (Andor). The second microscope was equipped with CSU-X1 spinning disk (Yokugawa), a MLC400B monolithic laser combiner (Agilent) with 405, 488, 561, and 640-nm laser lines, a 60x 1.4 NA Plan Apo objective (Nikon), and a Clara interline CCD camera. All microscopes were controlled using Nikon Elements.

For experiments where Bem1 was reactivated following 8 h of deactivation, images were acquired every 5 or 10 min. During imaging, a 725-nm longpass filter (FSQ-RG9, Newport) was placed in the transmitted light path (gently resting on top of the condenser). The shutter for the transmitted light source was then left open for the entire duration of the experiment (even when images were not actively being acquired), such that cells were constantly illuminated with IR light, ensuring that Bem1 remained ‘activated’. However, the shutter was briefly closed during acquisition of Cdc10-GFP images to reduce background.

### Image Analysis

Cell volumes for yeast undergoing isotropic growth (i.e., **Figs. 2B, 2D, 2F, and S1**) were measured using custom Matlab code (available upon request) where a Hough transform was applied to bright-field images to identify cell boundaries. Cell radii were determined by calculating the geometric mean of the major and minor axes of the resulting ellipse. “cell diameter fold-change” (e.g., **Fig. 2B**) was determined by calculating the ratio of the cell diameter at each timepoint to the initial diameter of the cell at “0 h”. A small fraction of cells did not arrest bud growth following Bem1 or Cdk1 inactivation and these were omitted from analysis.

For cells growing and dividing following Bem1/Cdk1 reactivation (i.e., **Figs. 3, 4, and S2**) cell volumes were calculated by manually fitting ellipses to the cell boundaries with a single focal plane passing through the center of the cell. The major and minor axes of the ellipses were then determined, and cell volumes were calculated by solving for the volume of a prolate spheroid. For analysis of bud ‘sizer’ and bud ‘timer’ mechanisms (**Fig. 4B-C**), we included in our analysis mother cells that had previously been Bem1- or Cdk1-deactivated. These cells were omitted from our analysis of total volume addition over the course of an entire cell cycle (**Fig. 3**), given that we had strongly perturbed their growth during G1 following Bem1 or Cdk1 disruption.

For the experiments depicted in **Fig. 3D-E**, all Bem1- and Cdk1-disrupted cells were from the same generation: daughters produced by giant mothers following Bem1 or Cdk1 reactivation. “Unperturbed” cells indicate optoBem1 cells that were grown in the absence of PCB (i.e. Bem1 activity was not light-responsive). Cells were imaged and analyzed as depicted in **Fig. 3B-C**. “volume at birth” for daughters was measured when the septin ring separating the mother and daughter disappeared (e.g., **Fig. 3C**, ‘00:15’, blue arrowhead). “G1 volume change” is the difference between volume measured when bud emergence occurred in newly-born daughters (e.g., **Fig. 3C**, ‘00:45’, yellow arrowhead), and “volume at birth”. “volume at division” was measured when daughters completed a subsequent round of cytokinesis (e.g., **Fig. 3C,** ‘02:45’, blue arrowhead) and included both the original daughter and her newly-formed bud. “interdivision volume change” is the difference between “volume at division” and “volume at birth”. A small fraction of both optoBem1 and *cdk1-ts* giant cells failed to bud following reactivation of Bem1 or Cdk1 and these were omitted from analysis. All cells quantified were obtained from two independent experiments, unless stated otherwise in the figure legends.

### Quantification of protein levels

To determine cellular protein levels in growing yeast, we quantified mCherry fluorescence in our cells, which expressed *BEM1-mCherry-PIF* under the endogenous *BEM1* promoter and *PhyB-mCherry-CAAX* under the *ADH1* promoter, by spinning-disc confocal microscopy. Isotropic growth was induced as described in the “Preparation of yeast for optogenetic experiments” section. Cells were imaged 6 h following optogenetic-based Bem1 deactivation. Z-stacks containing 51 slices with 0.5 µm spacing with the shutter remaining open between each z-step were then collected. Sum projections were created from the entire stack, corrected for uneven illumination, and processed to remove background. Fluorescence intensity was measured from whole-cell regions-of-interest using ImageJ. Cell volume was approximated by measuring the major and minor axis of each cell and using these values to solve for the volume of a prolate spheroid.

### Flow cytometry to measure ploidy

PCB-treated cells were placed in clear culture tubes and incubated at room temperature while undergoing constant illumination with a red LED panel. Samples (200 µL each) were collected at 0, 3, and 6 h following illumination with red light. The remainder of the culture was then placed under an IR LED (740-nm, Lightspeed Technologies) for an additional 12 h, after which a final sample was collected. Upon collection, cells were immediately treated with 475 µL 100% ethanol and fixed for 1 h at room temperature. Cells were then pelleted, washed once with 50 mM sodium citrate (pH = 7.2), and then resuspended in 500 µL 50 mM sodium citrate (pH = 7.2) to which RNaseA (0.25 mg/mL) had been added. Following an incubation for 1 h at 37 °C, 50 µL proteinase K was added to a final concentration of 2 mg/mL and cells were incubated at 50 °C for an additional hour. SYTOX green was then added to each sample to a final concentration of 2 µM, and cells were incubated for 1 h at room temperature. In parallel, control samples were prepared containing exponentially growing cultures of yeast treated with α-factor (10 µg/mL) for for ∼3 h prior to fixation and staining. Ploidy was determined by measuring the fluorescence intensity of SYTOX green staining by flow cytometry using a FACSAria III (BD Biosciences) and normalizing to the α-factor-treated samples, which have a ploidy of 1N. For each timepoint for each independent experiment, 50,000 cells were measured. Analysis of flow cytometry data was performed using FlowJo and Flowing Software (http://flowingsoftware.btk.fi).

### Calculation of doubling times

Prior to each experiment, cells were grown overnight to saturation in synthetic complete media at 25 °C. Cells were then diluted to an OD600 of 0.5-1.0 in fresh media and incubated while shaking in a Biotek H4 Plate Reader. The OD600 for each culture was measured every 5 min for 14-16 h, by which point growth curves reached saturation. Doubling times were then determined by plotting log2(OD600) vs. time and calculating the slope over the linear portion of the growth curve. Each point indicates an independent experiment.

## ACKNOWLEDGEMENTS

We thank Anna Payne-Tobin Jost for helpful discussions, Alba Diz-Muñoz for experimental assistance, Fred Chang for critical reading of the manuscript and Nairi Hartooni for reagents. We also thank the organizers of the 2015 MBL Physiology course, Wallace Marshall, Jennifer Lippincott-Schwartz, and Rob Phillips, as well as all the course staff for creating the well-run, intellectually-stimulating environment from which this project grew. Support for this work was provided by a Post-Physiology Course Award from the Marine Biological Laboratory to C.A.H.A., the Thomas B. Grave and Elizabeth F. Grave Scholarship to F.D., and the National Institutes of Health [GM118167 to O.D.W., DP2EB024247 to J.E.T., and T32HL773125 to B.R.G.].

## AUTHOR CONTRIBUTIONS

Conceptualization, all authors; Software, F.D. and J.E.T.; Investigation, C.A.H.A, F.D., J.E.T., and B.R.G.; Writing – original draft, B.R.G.; Writing – review and editing, all authors.

